# The Mehler reaction site is the Phylloquinone within Photosystem I

**DOI:** 10.1101/2020.08.13.249367

**Authors:** Marina Kozuleva, Anastasia Petrova, Yuval Milrad, Alexey Semenov, Boris Ivanov, Kevin E. Redding, Iftach Yacoby

## Abstract

Photosynthesis is a vital process, responsible for fixing carbon dioxide, and producing most of the organic matter on the planet. However, photosynthesis has some inherent limitations in utilizing solar energy. Up to a third of the energy absorbed is lost in the reduction of O_2_ to produce the superoxide radical (O_2_^•−^), which occurs principally within photosystem I (PSI) *via* the Mehler reaction. Strikingly, the precise location as well as the evolutionary role of the reaction have long been a matter of debate. For decades, O_2_ reduction was assumed to take place solely in the distal iron-sulfur clusters of PSI rather than within the two asymmetrical cofactor branches. Here we demonstrate that under high irradiance, O_2_ photoreduction by PSI takes place at the phylloquinone of one of the branches (the A-branch). This conclusion derives from the light dependency of the O_2_ photoreduction rate constant, and from the high rates of O_2_ photoreduction in PSI complexes lacking iron-sulfur clusters and in a mutant PSI, in which the lifetime of this phyllosemiquinone state is extended 100-fold. On these grounds, we suggest that the Mehler reaction serves as a release valve, functioning only when needed, under conditions where both the distal iron-sulfur clusters of PSI and the mobile ferredoxin pool are over reduced.

**SIGNIFICANCE STATEMENT:** Photosynthesis is the process responsible for the oxygenation of the ancient anoxic atmosphere, and the transformation of inorganic carbon to most of the organic matter on Earth. However, it is less commonly appreciated that the appearance of oxygen in the atmosphere led to the unavoidable opposite process in which oxygen is consumed, thereby producing deleterious oxygen radicals such as the superoxide radical. For almost half a decade, the location of the main site of superoxide radical production in chloroplasts has been a matter of debate. We now provide conclusive evidence that it is located in the phylloquinones(s) within photosystem I.

## INTRODUCTION

Under high irradiance, oxygenic phototrophs can use molecular oxygen (O_2_) as an alternative sink for surplus electrons within the photosynthetic apparatus (1). Unfortunately, uncontrolled leakage of electrons to O_2_ could decrease the quantum yield of photosynthesis and produce reactive oxygen radicals. As a result, the photosynthetic apparatus appears to have evolved strong regulation of such reactions in an aerobic environment (2).

The production of hydrogen peroxide (H_2_O_2_) in spinach thylakoid membranes under illumination was first reported by Alan Mehler (3). Later, the superoxide radical (O_2_^•−^) was shown to be the primary product of this reaction (4). Nowadays, O_2_ photoreduction to O_2_^•−^ by a photosynthetic electron transfer (ET) chain is referred to as the true Mehler reaction, in order to distinguish it from a Mehler-like reaction (5, 6). Although some evidence indicates that O_2_^•−^ production can take place in various locations (2, 7– 9), photosystem I (PSI) seems to be the major site of this process (10, 11).

The PSI complex mediates a light-driven charge separation that is accompanied by plastocyanin (Pc) oxidation and ferredoxin (Fd) reduction (Fig. 1A). Two large membrane subunits, PsaA and PsaB together with a smaller extrinsic subunit PsaC coordinate the ET cofactors comprising six chlorophyll *a* (Chl) molecules, two phylloquinone (PhQ) molecules, and three [4Fe-4S] clusters. With the exception of the [4Fe-4S] clusters, all the other cofactors are organized within two asymmetric branches (named A- and B-). Both the A- and B-branches are active (12). One of the most critical differences between them is related to the redox properties of PhQs, where a higher potential quinone is located in the A-branch, while the B-branch has a lower potential quinone (Fig. 1A). Notably, the environment influences the properties of PhQ at the A_1_-sites, so that the E_m_ and lifetimes of the phyllosemiquinones (PhQ^•−^) are modulated *via* the identity of nearby amino acids. The most drastic effect published so far was due to the replacement of PsaA-Phe_689_ with Asn (PsaA-F_689_N mutant), which resulted in a ∼2 orders of magnitude longer lifetime of PhQ^•−^ at the A-branch (PhQ_A_^•−^) (from ∼0.25 µs to 17 µs) (13), probably as a result of a ∼125-mV upshift of the PhQ_A_/PhQ_A_^•−^ reduction potential (Fig. 1B).

**Fig. 1.**
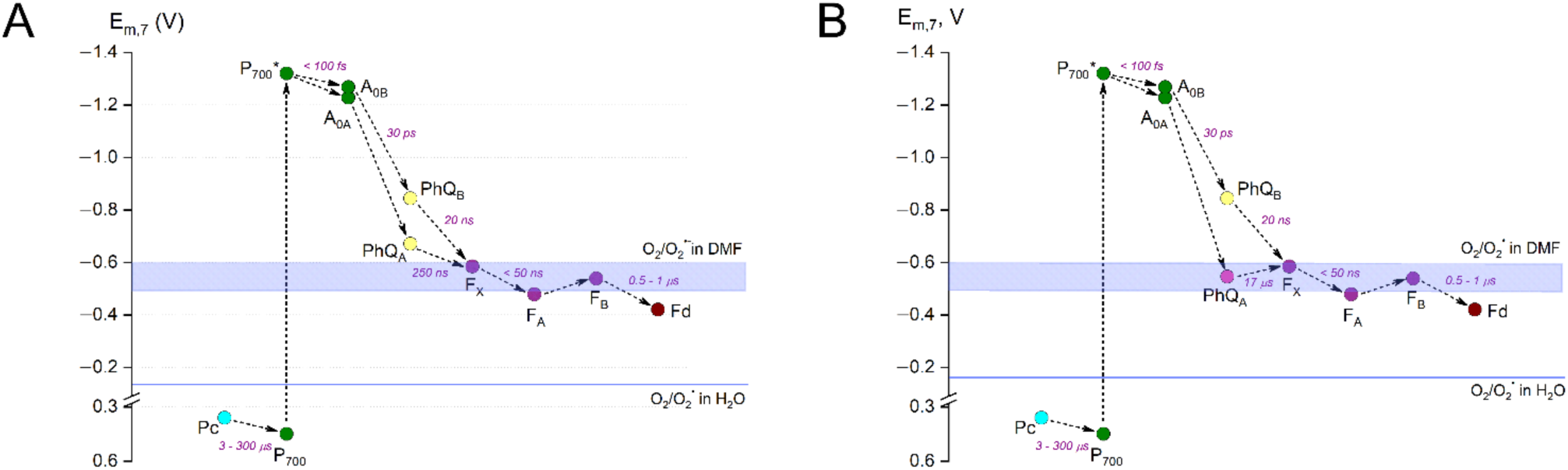
Diagram of electron transfer in Photosystem I of wild type (A) and PsaA-F_689_N (B) with lifetimes and E_m_ of cofactors according to (14); blue lines represent E_m_ of O_2_/O_2_^•−^ in water (−160 mV) and dimethylformamide (−500 - −600 mV) (15). Modified from (2).

Oxygen photoreduction in PSI is generally thought to occur in the F_A_/F_B_ clusters. This common belief stems from the fact that, in isolated thylakoid membranes, F_A_/F_B_ are the terminal cofactors, and they reduce O_2_ under steady-state illumination in the absence of Fd (11, 16–18). However, the primary site of O_2_ photoreduction in thylakoid membranes was shown to be the heterodimer PsaA/PsaB, rather than PsaC carrying F_A_/F_B_ clusters (11). A contribution of PhQs to the Mehler reaction was first suggested by (19) and a first experimental evidence for the involvement of PhQs in O_2_ photoreduction under steady-state illumination was provided by (20). However, a contribution of PhQ at the A- and B-branches as well as contribution of [4Fe-4S] clusters, especially the intermediate F_X_ cluster, are still open questions. The mechanisms underlying the O_2_^•−^ production within PSI occurring concomitantly with electron transfer to Fd and then to NADP^+^ and with varying irradiance are still unclear.

The aim of the present study was to identify where and how photoreduction of O_2_ occurs within PSI. Towards that goal, we measured the apparent second-order rate constant of O_2_ photoreduction by PSI complexes (*k*_*2*_) as a function of irradiance. We investigated intact fully mature PSI complexes isolated from a model organism, green alga *Chlamydomonas reinhardtii*, and PSI complexes lacking either the F_A_/F_B_ (F_X_-core), or all three [4Fe-4S] clusters (A_1_-core). In addition, we investigated PSI complexes harboring the PsaA-F_689_N mutation, which results in a long-lived PhQ_A_^•−^ (13). Our results strongly support the conclusion that PhQs, especially PhQ_A_, are the major site of O_2_ photoreduction within PSI under high irradiance.

## RESULTS

### Under continuous illumination, O_2_ photoreduction is the rate-limiting step of electron transfer

Identification of the sites of O_2_ photoreduction within PSI under steady-state illumination requires the ET to O_2_ to be the rate-limiting step of total ET. This can be accomplished by providing an efficient electron donor to P_700_^+^. Here, we used excess Pc to reduce P_700_^+^ in isolated fully mature intact PSI complexes. In order to verify the Pc concentration, we used the approach described in (20) for cyanobacterial PSI complexes and *N,N,N′,N′*-tetramethyl-*p*-phenylenediamine (TMPD) as an electron donor to P_700_^+^. Since increasing the concentrations of Pc in this model did not affect the O_2_ photoreduction rate (Fig. 2A), we conclude that the O_2_ photoreduction is indeed the rate-limiting step of the overall reaction. The same result was observed in our system when Pc was replaced by TMPD (Fig. S1). Although the rate constant for TMPD (3.0× 10^4^ M^− 1^ s^− 1^ (21)) is ∼ four orders of magnitude lower than that of Pc, the rates of O_2_ photoreduction were the same. Neither did increasing the concentration of electron donor affect the rate of O_2_ photoreduction when all the [4Fe-4S] clusters were removed (Fig. S2), although the basal rate was higher (see further). In contrast, when MV, an efficient redox-mediator from PSI to O_2_, was added to intact PSI complexes, the rate of O_2_ photoreduction did increase with increasing Pc concentration (Fig. 2A). Thus, we can conclude that the electron donation to P_700_^+^ from Pc/TMPD was the rate-limiting step only in the presence of efficient electron acceptors such as MV.

**Fig. 2.**
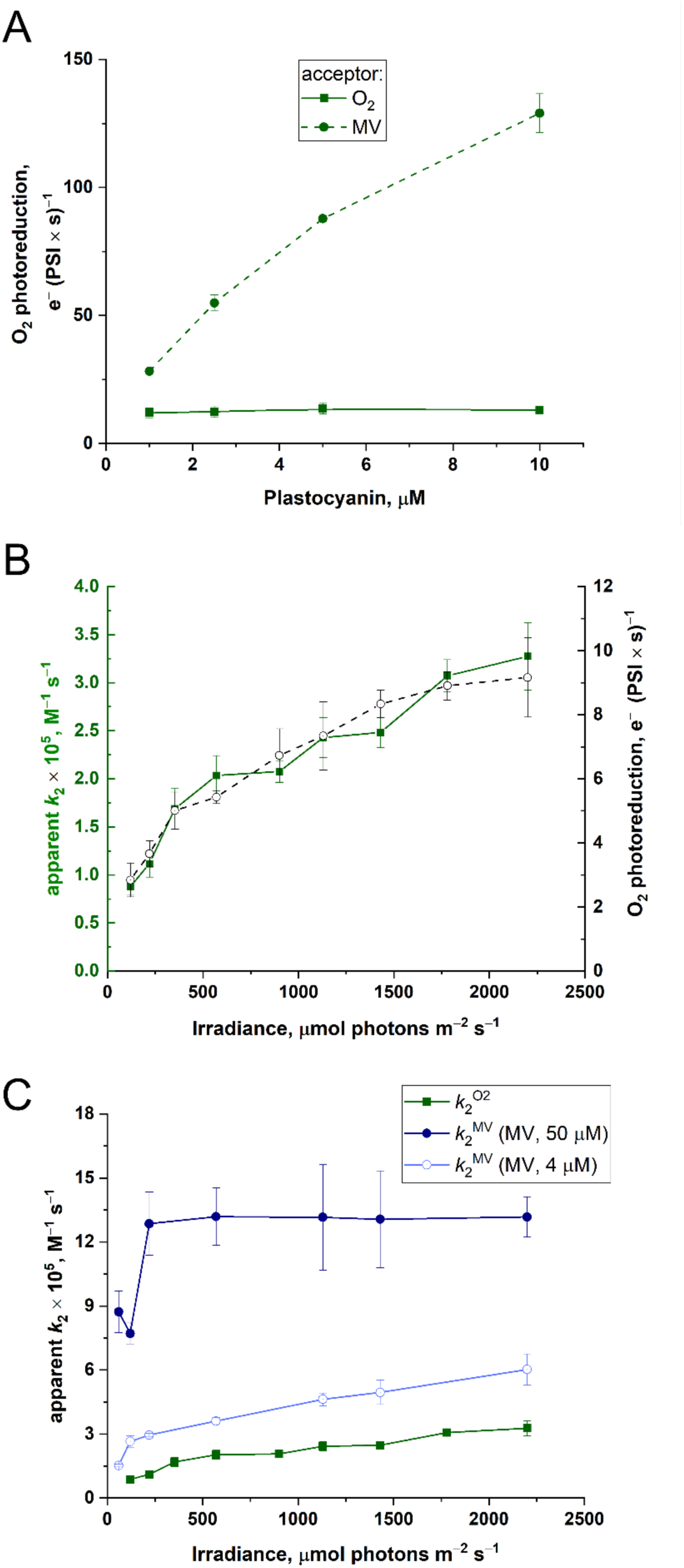
Effect of plastocyanin concentration on the O_2_ photoreduction rate in the absence or presence of MV at 50 µM (A) and effect of irradiance on the apparent second order rate constant of O_2_ photoreduction by PSI in the absence of MV (B) and in the presence of MV at 4 µM (light blue line) and 50 µM (dark blue line) (C); Panel B, the O_2_ photoreduction rate (dashed lines) is shown for comparison; Panel C, the intact PSI (green line) is shown for comparison. The experiments conditions were: Mature wild type PSI complexes, 10 nM (B and C) or 15 nM (A); Asc, 10 mM; catalase, 500 U ml^−1^ (B and C); Pc, varied (A) or 5 µM (B and C); initial O_2_ concentration, 10-30 µM for measurement of apparent *k*_2_ (B and C) and 250 µM for measurement of O_2_ photoreduction rate (A and black dashed lines on panel B); irradiance, 2200 µmol photons m^−2^ s^−1^ (A). Values are mean for 7-10 replicates for *k*_2_ and 3-4 replicates for rates; ±S_E_.

Our working hypothesis is that under steady-state illumination, there are multiple oxygen photoreduction sites in PSI, each catalyzing a mechanistically independent reaction associated with a discrete rate constant. The accumulation of electrons on these sites changes in response to irradiance. In other words, each site reaches maximal activity at a different specific irradiance. Therefore, if the apparent second-order rate constant of O_2_ photoreduction by PSI (*k*_2_) is not the same at all light levels, but is rather a function of irradiance, it means that multiple oxygen photoreduction sites are involved. A physical model to explain this behavior would be to postulate that a site must be reduced in order to be able to transfer an electron to O_2_. If there are few electron acceptors, electrons become “stuck” in the system, with the terminal sites filling up first. Hence, electrons start to accumulate at the F_A_/F_B_ centers under low irradiance and then, with increasing irradiance, they begin to occupy the F_X_ site and then A_1_-sites.

### Determination of the second-order rate constant of O_2_ photoreduction in PSI

The exact rate constant of the Mehler reaction is not currently known and estimates for the kinetic parameters for O_2_ photoreduction by PSI range from ∼10^3^ M^−1^ s^−1^ (22) to 10^7^ M^−1^ s^−1^ (23). The mid value (7 × 10^4^ M^−1^ × s^−1^) was obtained by kinetic modeling of charge recombination kinetics observed after a light flash (24). We measured the apparent *k*_2_ for intact fully mature PSI complexes purified from the wild type (WT) strain (PSI^WT^), under conditions where O_2_ was the principal electron acceptor. In this case, *k*_2_ was denoted as *k*_2_^O2^. The apparent *k*_2_^O2^ varied slightly between different PSI preparations and was estimated to be ∼3.5 ×10^5^ M^−1^ s^−1^ at the highest irradiance tested (2200 µmol photons m^−2^ s^−1^). Fig. 2B shows the dependency of the apparent *k*_2_^O2^ on irradiance as measured in a typical PSI preparation. The apparent *k*_2_^O2^ clearly increased with irradiance up to 2200 µmol photons m^−2^ s^−1^. The O_2_ photoreduction rate measured at atmospheric pressure of O_2_ as a function of irradiance resembles that of *k*_2_^O2^ (Fig. 2B, dashed line), as expected.

### Saturating concentrations of Methyl Viologen (MV) *de facto* produce a single O_2_ reducing site

As F_A_/F_B_ are oxidized completely by an excess of MV, the effect of irradiance was examined in the presence of MV at saturating concentration (50 µM). In this case, the measured constant was denoted as *k*_2_^MV^ since MV was the principal electron acceptor from PSI and was constantly recycled by reduction of O_2_. Indeed, the apparent *k*_2_ and O_2_ uptake rates were higher in the presence of MV (Fig. 2A and 2C), reflecting higher PSI turnover. While direct O_2_ photoreduction by PSI cofactors can take place at the same time as MV reduction, it will be largely suppressed at saturating MV concentrations. In contrast to behavior of *k*_2_^O2^, the values of the apparent *k*_2_^MV^ reached a plateau at 220 µmol photons m^−2^ s^−1^ (Fig. 2C, dark blue circles), demonstrating that in the presence of high concentrations of MV, there is *de facto* a single O_2_ photoreduction site. However, when MV was added at a 12-fold lower concentration (4 µM), nearing the K_m_(MV) value determined in separate experiments (Fig. S3), *k*_2_^MV^ increased with increasing irradiance and did not reach a plateau (Fig. 2C, light blue circles), *i*.*e*., the behavior of *k*_2_^MV^ at 4 µM MV was more similar to that of *k*_2_^O2^. Since at concentrations around the Km, MV would account for only about half of the maximal rate, we conclude that electrons are still accumulating at intermediate PSI cofactors and are available for direct reduction of O_2_. In order to negate the possibility that the saturation behavior of *k*_2_^MV^ at 50 µM MV is due to limitations in re-reduction of P_700_^+^, we replaced Pc by TMPD, a less efficient electron donor than Pc. In the presence of TMPD, the apparent *k*_2_^MV^ was lower but still reached a plateau at the same irradiance (Fig. S4), confirming that the irradiance dependence of the apparent *k*_2_^MV^ is not influenced by processes taking place at the donor side of PSI.

### Stripping PSI of its [4Fe-4S] clusters enhance O_2_ photoreduction

The hypothesis that there are multiple sites within PSI for O_2_ photoreduction can be tested by removal of the terminal cofactors. This should remove the competition for electrons and favor reduction of O_2_ by upstream cofactors (*i*.*e*., F_X_, PhQ_A_, and PhQ_B_). To test the role of F_X_ and PhQ in O_2_ photoreduction, we sequentially stripped PSI of the [4Fe-4S] clusters. Initially, the subunit PsaC containing the F_A_/F_B_ clusters was removed, resulting in a PSI complex, in which the terminal cofactor is F_X_ (F_X_-core complexes). The behavior of the apparent *k*_2_^O2^ in the F_X_-core complexes was rather similar to that of the intact complexes (Fig. 3A, yellow circles *vs*. green squares), especially at high irradiance, with no stimulation of O_2_ photoreduction observed. This result suggests that F_X_ plays a minor role in O_2_ photoreduction in PSI. Removal of F_A_/F_B_ actually decreased O_2_ reduction at low irradiances. This result supports the contention that F_A_/F_B_ plays a role in O_2_ reduction in low light. As the next stage, we removed F_X_, leaving the bound PhQ molecules at the A_1_-sites as the terminal cofactors (A_1_-core complexes). In contrast to the results obtained with the F_X_-core, A_1_-core complexes showed markedly higher rates of O_2_ photoreduction at irradiance above 350 µmol photons m^−2^ s^−1^ (Fig. 3A, blue triangles vs. green squares). Interestingly, O_2_ photoreduction in the A_1_-core complexes in low light is faster than in F_X_-core complexes. This indicates that F_X_ is a poor reductant of O_2_. Taking together, these results demonstrate that none of the [4Fe-4S] clusters is essential for photoreduction of O_2_ by PSI. Furthermore, they indicate that the PhQs are the moieties responsible for O_2_ photoreduction in intact PSI in high light.

**Fig. 3.**
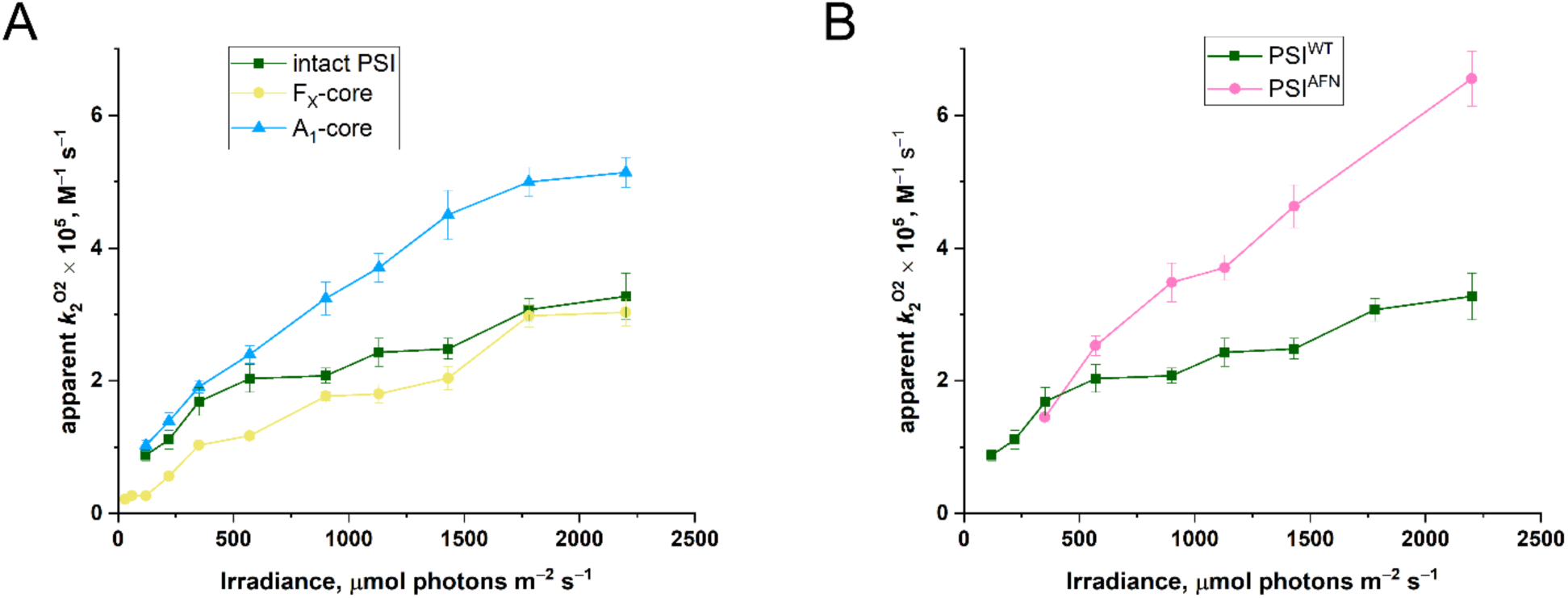
Effect of irradiance on the apparent rate constant of O_2_ photoreduction by F_X_- and A_1_-core complexes (A) and by the PsaA-F_689_N mutant (B). Intact wild type PSI (green line) is shown for comparison. The experiments conditions were: F_X_-core, A_1_-core complexes, PSI^AFN^, 10 nM; TMPD, 1 mM; Asc, 10 mM; catalase, 500 U ml^−1^; initial O_2_ concentration, 10-30 µM. Values are mean for 7-10 replicates; ±S_E_.

### A mutant PSI^AFN^, in which PhQA^•−^ has a longer lifetime, demonstrates dramatically enhanced O_2_ photoreduction

One might argue that the results with the A_1_-core complexes were merely due to unspecified damage caused by the removal of the [4Fe-4S] clusters. In order to test our hypothesis in intact PSI particles, we made use of the PsaA-F_689_N mutant, in which PhQ_A_^•−^ has a lifetime about two orders of magnitude longer than in the WT (13). Slowing down ET from PhQ_A_ should increase the chances that O_2_ can catch electrons from PhQ_A_^•−^ *en route* to downstream cofactors, obviating the need to remove them. The behavior of charge recombination kinetics in PSI complexes from this mutant (PSI^AFN^) more closely resembled that of WT A_1_-core complexes than intact PSI^WT^ (Fig. S5, Petrova, in preparation) showing a diminished population with the reduced F_A_/F_B_. Here, we used mature PSI^AFN^ complexes capable of reducing NADP^+^ in the presence of Fd and FNR under steady-state illumination (Table S1). However, the rates of NADP^+^ reduction in PSI^AFN^ complexes were 2-5 times lower than in PSI^WT^. Our hypothesis predicts faster rates of O_2_ reduction in the PsaA-F_689_N mutant than in intact PSI^WT^ complexes. Indeed, at intensities above 350 µmol photons m^−2^ s^−1^, the values of the apparent *k*_2_^O2^ in PSI^AFN^ were markedly higher than those of intact PSI^WT^ (Fig. 3B, pink circles vs. green squares) and the curve resembled that of WT A_1_-core complexes (Fig. 3A, blue triangles) more than WT intact PSI.

### O_2_ photoreduction does not compete with NADP^+^ reduction

One might question the relevance of these *in vitro* results to the situation *in vivo*, where electron acceptors such as Fd are available to PSI. In order to assess the impact of such acceptors upon O_2_ photoreduction by PSI, we repeated the experiment with the addition of Fd, FNR, and NADP^+^, *i*.*e*. under conditions, which simulate the situation *in vivo*. The presence of FNR and excess NADP^+^ provided an efficient electron sink for Fd, as evidenced by the measured NADP^+^ photoreduction rate (Fig. 4B). Photoreduction of O_2_ by components of the thylakoid membrane has been shown to occur concomitantly to reduction of NADP^+^ (18). Furthermore, Fd also contributes to O_2_ reduction, especially in the absence of NADP^+^. Hence, the apparent *k*_2_ in the presence of Fd (*k*_2_^Fd^) should be the sum of *k*_2_ for ET from PSI to O_2_ either directly or *via* Fd. Indeed, in the presence of Fd alone, the values of the apparent *k*_2_^Fd^ were higher than *k*_2_^O2^ (Fig. 4A), suggesting a strong contribution of Fd to the total O_2_ reduction under such conditions. However, in the presence of FNR and NADP^+^, we observed similar rates of O_2_ photoreduction at high irradiance as in the absence of Fd/FNR/NADP^+^ (Fig. 4A, violet circles *vs*. green squares). Hence, electron flow to NADP^+^ probably decreases the Fd-dependent O_2_ reduction by inhibiting the accumulation of reduced Fd. Interestingly, in contrast to the situation at high irradiance, the presence of Fd/FNR/NADP^+^ slows down O_2_ photoreduction at low irradiance, indicating that NADP^+^ reduction also decreases the accumulation of electrons on [4Fe-4S] clusters. The similarity in behavior of *k*_2_^Fd^ in the presence of FNR/NADP^+^ and *k*_2_^O2^ at high irradiance strongly suggests that PhQs at the A1-sites represent the major site of O_2_ photoreduction in PSI *in vivo* under high light.

**Fig. 4.**
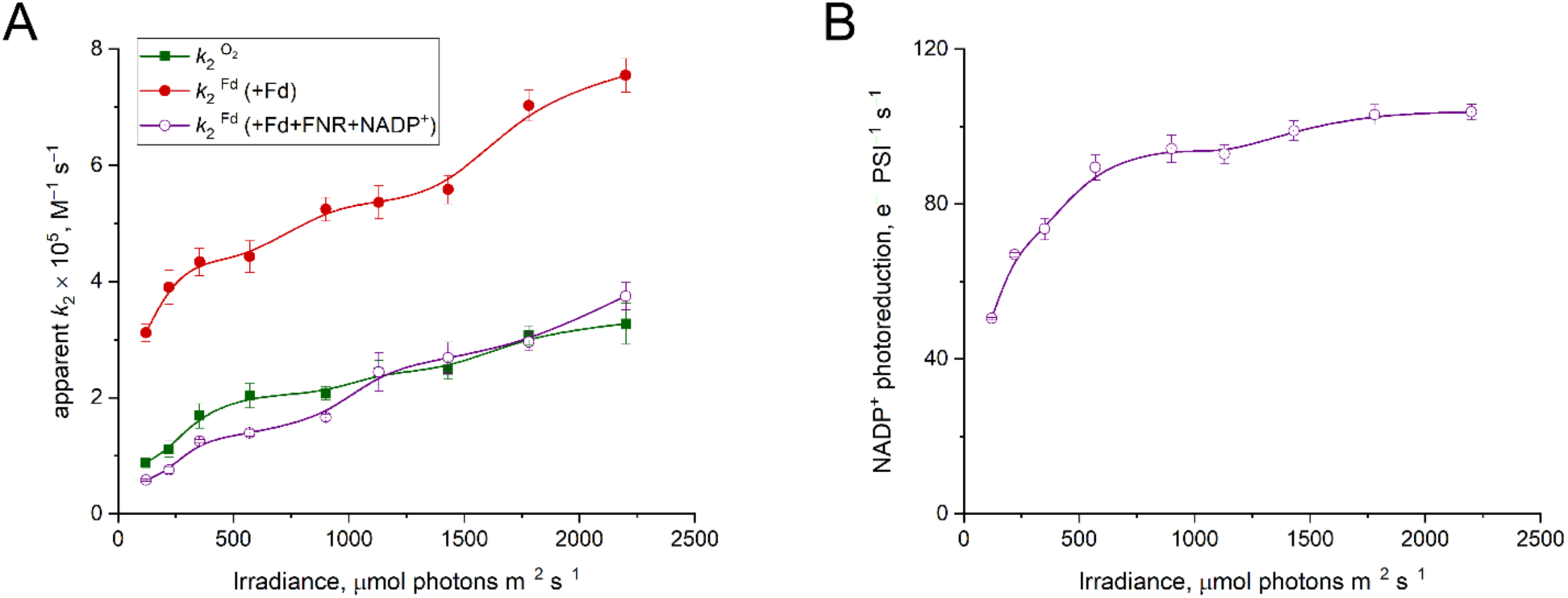
Effect of irradiance on the apparent rate constant of O_2_ photoreduction in the presence of Fd and Fd + FNR + NADP^+^ (A) and on the rate of NADP^+^ reduction (B); the intact PSI (green line) is shown for comparison. The experiments conditions were: Mature wild type PSI complexes, 10 nM; Pc, 5 µM; Asc, 10 mM; catalase, 500 U/ml; where indicated Fd, 5 µM; FNR, 200 nM; NADP^+^, 1 mM; initial O_2_ concentration, 10-30 µM. Values are mean for 7-10 replicates for *k*_2_ and 5 replicates for rates; ±S_E_.

## DISCUSSION

In this study, we measured the rates of O_2_ photoreduction by various PSI complexes and demonstrated that the apparent rate constant of this reaction depends on irradiance (Fig. 2A). This observation could be explained by either *i*) the presence of a single O_2_ reduction site (*i*.*e*. F_A_/F_B_), that attains full activity under high irradiance, or *ii*) the presence of multiple O_2_ reduction sites within PSI, which are activated under different light regimes. Our data are much more consistent with the latter model.

### F_A_/F_B_ is neither the sole nor the major O_2_ photoreduction site within PSI

The observation that the removal of F_A_/F_B_ slows the O_2_ reduction rate at low irradiances indicates their responsibility for the process under low light (Fig. 3A). The results obtained following MV addition to the intact PSI complexes (Fig. 2C) demonstrate that there is unlikely to be a single O_2_ photoreduction site in PSI. With saturating concentrations of MV, which *de facto* simulate the behavior of a single site of O_2_ reduction, *k*_2_^MV^ reached a plateau at an irradiance of 220 µmol photons m^−2^ s^−1^. Since in intact PSI, MV accepts electrons exclusively from F_A_/F_B_, and not from intermediate cofactors, this value represents the optimal irradiance for activating F_A_/F_B_. In other words, if *k*_2_ increases any further in the absence of MV, it must indicate reduction of O_2_ on upstream cofactor(s). Moreover, the contribution of upstream cofactor(s) to O_2_ reduction must depend on the rate of electron withdrawal from PSI by available acceptors, since a *k*_2_^MV^ plateau was no longer observed when lower concentrations of MV were provided (Fig. 2C) and the behavior was similar to that of *k*_2_^O2^ at higher irradiances.

### Under high irradiance, PhQ_A_ is the major site of O_2_ photoreduction in PSI

In order to localize the site(s) of O_2_ photoreduction in PSI, we removed the [4Fe-4S] clusters sequentially. Stripping PSI of F_A_/F_B_ lowered O_2_ photoreduction rates at low irradiances but had little effect at high irradiances (Fig. 3A). The additional removal of F_X_ resulted in a significant increase in *k*_2_^O2^ values at irradiances exceeding 350 µmol photons m^−2^ s^−1^. These results demonstrate that PhQs at the A_1_-sites, and not F_X_, are the major sites of O_2_ reduction in intact PSI under high irradiance. The same conclusion regarding the contribution of PhQs to O_2_ photoreduction was reached independently *via* experiments using the mature PSI particles from PsaA-F_689_N mutant, in which the lifetime of PhQ_A^•−^_ is increased from ∼0.2 µs to ∼17 µs (13). At irradiances higher than 350 µmol photons m^−2^ s^−1^, the *k*_2_^O2^ values for PSI^AFN^ complexes were significantly higher than those measured for the PSI^WT^ complexes (Fig. 3B). Our interpretation is that the longer lifetime of PhQA^•−^ increases the chances of this semiquinone reacting with O_2_. Note that the competing hypothesis that [4Fe-4S] cluster(s) are the primary site(s) of O_2_ reduction at high irradiance, would have predicted the opposite result, as the population with reduced clusters is diminished in this mutant (Fig. S5, Petrova, in preparation). This is consistent with our observation of a 2-5 times lower rate of NADP^+^ photoreduction under steady-state illumination (Table S1) as well as a decrease in MV-driven O_2_ reduction rates (Fig. S7) by PSI^AFN^ than by PSI^WT^ complexes.

Santabarbara reported that the lifetime of PhQ_B_^•−^ in PsaA-F_689_N mutant is the same as in WT (13) and is ∼one order of magnitude lower than the lifetime of PhQ_A_^•−^ in WT. This decreases the chances of PhQ_B_^•−^ being an efficient contributor to O_2_ photoreduction in intact PSI. Taking all these points into consideration, we hypothesize that the results presented in Fig. 3B, demonstrate that PhQ_A_^•−^ rather than PhQ_B_^•−^ is responsible for O_2_ photoreduction.

### O_2_ photoreduction does not compete with NADP^+^ reduction

Our data revealed that at high irradiance, the major contribution of PhQs within PSI to O_2_ photoreduction is valid even when Fd and NADP^+^ reduction takes place, *i*.*e*. under conditions simulating physiological conditions. Addition of Fd alone significantly increased the rates of O_2_ reduction (Fig. 4A). This partially imitates the situation *in vivo* when the stromal NADP^+^ pool is in the reduced state (*e*.*g*., due to a retarded Calvin cycle at limiting CO_2_). However, since we used Fd at much higher levels relative to PSI than present *in vivo* (500:1 ratio *vs*. 5:1 ratio in spinach chloroplast; (25)), care should be taken when extrapolating this result to the *in vivo* situation. It is likely that the addition of FNR and NADP^+^ minimized the Fd-dependent O_2_ reduction by decreasing the accumulation of reduced Fd. These observations are in good agreement with previously reported conclusions (18, 26). The results of our experiments indicated saturation of NADP^+^ photoreduction at high irradiances (Fig. 4B). This provides insights into the *in vivo* situation, since these are conditions, when the Mehler reaction plays a physiological role in protecting the photosynthetic apparatus from over-reduction of the ET chain and when the PhQs pass electrons to O_2_. This can suggest a crucial role for PhQs, and specifically PhQ_A_ (see above), in O_2_ photoreduction in PSI *in vivo*.

Here we used PSI complexes isolated from *C. reinhardtii*. The acceleration of H_2_O_2_ in the chloroplasts as well the increase in enzymatic antioxidant capacity was shown to occur when *C. reinhardtii* cells were subjected to high light (27). However, the physiological significance of the true Mehler reaction in angiosperms is much well recognized. Since the composition of the ET cofactors in PSI is highly conserved through all green lineages, O_2_ photoreduction by PhQs represents a universal mechanism of the Mehler reaction.

## MATERIALS AND METHODS

### Strains, growth, thylakoid membrane and PSI isolation

*C. reinhardtii* strains with His_6_-tag added to the N-terminus of the PsaA protein (PBC1 pKR152 for WT and PBC1 pKR411 for the strain bearing a mutation in codon 689 of *psaA* exon 3 (*psaA-3*)) were grown in TAP medium at 25 °C with constant air flow under continuous white light (∼70 µmol photons m^−2^ s^−1^). The culture was harvested, and thylakoid membranes and mature PSI complexes were purified as described in (28). The fraction corresponding to the mature PSI complexes was collected, concentrated, and stored at −80 °C. The maturity of the complexes was confirmed (Table S1).

### Expression and purification of recombinant proteins

The recombinant *C. reinhardtii* Pc, Fd and ferredoxin-NADP^+^ oxidoreductase (FNR) were heterologously expressed in *Escherichia coli* BL21 cells, and purified as described in (28) and (29).

### Determination of P_700_^+^

The concentration of photo-oxidized P_700_^+^ was determined by measuring light minus dark absorption changes at 698 nm (ε = 100 mM^−1^ × cm^−1^)(30). The assay medium contained 30 µg Chl ml^−1^ PSI, 20 mM NaCl, 5 mM MgCl2, 10 mM Asc, 10 nM Pc, 50 µM methyl viologen (MV), 0.03% β-DDM, 20 mM Tricine-NaOH (pH 7.5) was placed in spectrophotometer (Cary 50, Varian) and illuminated with saturating white light (Intralux 5000, Volpi).

### Measurement of O2 concentration changes using MIMS

Changes in O_2_ (the stable isotope ^16^O_2_) concentration were measured with a Membrane Inlet Mass Spectrometer (MIMS, QMS 200 M1; Pfeiffer Vacuum) according to (31); the data collection frequency was 1 s. Measurements were conducted under constant stirring at 21°C. The signal was calibrated by calculating the difference in values obtained in air saturated buffer (containing 250 µM O_2_) and in O_2_-depleted buffer after removal of O_2_ by glucose (10 mM), glucose oxidase (30 U ml^−1^), and catalase (500 U ml^−1^). A suspension of PSI complexes in buffer containing 20 mM NaCl, 5 mM MgCl_2_, 0.03% β-DDM, 20 mM Tricine-NaOH (pH 7.5), and electron donors and acceptors as indicated in figure legends, was placed in a sealed cuvette and illuminated by red light provided by a Dual-PAM (Walz GmbH).

In preliminary experiments, using the approach described in (20, 32), we confirmed that light-dependent O_2_ consumption resulted from direct ET from PSI cofactors to O_2_ (Table S2 and Table S3). The light-induced rate of O_2_ consumption (V_O2_) was calculated by subtraction of the dark slope from the light slope.

In experiments aimed for determination of *k*_2_, the O_2_ concentration in the reaction medium was decreased to 10-30 µM by N_2_ flushing of the headspace over the suspension in the sealed vessel, followed by transfer of the suspension by syringe to the air-depleted sealed cuvette containing the MIMS inlet probe and stir bar. It was necessary to restrict the O_2_ concentration because *k*_2_ can only be determined if the rate of the reaction depends on the concentration of O_2_ (Fig. S6). In these experiments, catalase (500 U ml^−1^) was added to prevent accumulation of H_2_O_2_, which is decomposed by [4Fe-4S] clusters to give a highly reactive hydroxyl radical capable of inhibiting PSI. In addition, accumulated H_2_^16^O_2_ molecules (mass 34) could potentially interfere with the ^16^O_2_ (mass 32) signal, and introduce errors. The rate constant was calculated as described in Appendix.

### Preparation of F_X_-core and A_1_-core

The F_X_-core complexes were obtained by incubating PSI complexes in buffer containing 6.8 M urea (33). The A_1_-core complexes were obtained by incubating F_X_-core complexes in buffer containing 3.4 M urea and 5 mM K_3_[Fe(CN)_6_]. The treatment efficiency was followed by monitoring the acceleration of charge recombination kinetics from ca. 50 ms in the intact PSI complexes to 1 ms and 10 µs in F_X_-core and A_1_-core respectively (Fig. S5).

### NADPH measurement

Light induced NADPH production was detected by a Dual-PAM equipped with DUAL-ENADPH and DUAL-DNADPH modules. The signal was calibrated by adding a known concentration of NADPH.

### Chlorophyll extraction and measurement

Chl concentration was determined spectrophotometrically after extraction in 96% ethanol according to (34).

## Acknowledgements

This work was supported by the ISF (Israel Science foundation) 1646/16, BSF (US Israel binational science foundation) 201666, The Ministry of Science and Higher Education of the Russian Federation, State Scientific Program, no. AAAA-A17-117030110135-1; the investigation of F_X_-core and A_1_-core complexes was supported by the RSF (Russian Science Foundation) 17-04-01323. M.K. was supported by the Fulbright Visiting Scholar Program. M.K. is thankful to Dr. Syed Lal Badshah, Mr. Oren Ben-Zvi, and Dr. Pini Marco for help with protein purifications, Patricia Baker for strain construction and help with alga culture maintenance, and to Dr. Ilya Naydov and Dr. Maria Borisova-Mubarakshina for valuable discussion.

