## Supplement for "The Mehler reaction site is the Phylloquinone within Photosystem I"

**This PDF file includes:**

Figures S1 to S7
Tables S1 to S3

**Table S1.** Characterization of intact PSI complexes purified from WT and the mutant PsaA-$F_{689}$N.

| Type of complexes | | Chl *a*/*b* ratio | Chl/$P_{700}$ | $NADP^+$ reduction rate, ($NADP^+$ ($P_{700}$ s)$^{-1}$) |
|---|---|---|---|---|
| The wild type | Preparation I | 5.6 | 223 | 101.8 |
| | Preparation II | 5.6 | 284 | 52 |
| | Preparation III | 5.0 | 328 | 57.2 |
| PsaA-$F_{689}$N | Preparation I | 5.3 | 327 | 20.8 |
| | Preparation II | n.d. | 250 | 21.8 |

The maturity of the PSI complexes was confirmed according to (Marco et al., 2018). The Chl *a*/ Chl *b* (*a/b*) ratio was 5.4 ± 0.1 indicating the presence of Chl *b*-containing Lhca proteins, which are lacking in immature PSI complexes. The Chl/$P_{700}$ ratio was 282 ± 21, indicating the presence of Lhca proteins. Immature PSI complexes are characterized by a low Chl/$P_{700}$ ratio (ca. 100) since ~100 Chl *a* molecules are bound to the PsaA/PsaB heterodimer per 1 $P_{700}$. All preparations of PSI used efficiently reduced $NADP^+$ at saturating light in the presence of Asc, Pc, Fd, FNR and $NADP^+$ with no artificial donors added. Addition of the artificial donor to $P_{700}^+$ DCPIP (50 µM) had no effect on the rate of $NADP^+$ reduction in studied complexes (not shown).

### Material and Methods

#### Calculation of the ratios Chl/$P_{700}$ and Chl *a/b*

The Chl/$P_{700}$ ratio was determined for each PSI preparation assuming all Chls are Chl *a*, MW = 893.5 g/mol. Chl *a/b* ratio was calculated according to (Lichtenthaler, 1987).

#### NADPH measurement

The maximal rate of NADPH production was detected as the change in absorbance at 340 nm ($\varepsilon = 6.27\ mM^{-1} \times cm^{-1}$) in PSI solubilized in buffer containing 20 mM NaCl, 5 mM $MgCl_2$, 0.03% β-DDM, 10 mM ascorbate, 10 µM Pc, 5 µM Fd, 200 nM FNR, 1 mM $NADP^+$, and 20 mM Tricine-NaOH (pH 7.5), and placed in a quartz cuvette. Values were measured with a Cary 50. The saturating white light was provided by an Intralux 5000, Volpi.

**Table S2.** Effect of superoxide dismutase and catalase on the rate of $O_2$ uptake in a suspension of fully mature wild type PSI complexes.

| | Rate of $O_2$ uptake, $O_2\ (P_{700} \times s)^{-1}$ | | Effect of enzyme addition |
|---|---|---|---|
| | no enzyme | + enzyme | |
| superoxide dismutase | 20.28 ± 2.41 | 9.78 ± 1.87 | ~ twofold decrease |
| catalase | 17.39 ± 3.51 | 8.74 ± 1.78 | ~ twofold decrease |

Mature wild type PSI complexes, 8 nM; Pc, 5 µM; Asc, 10 mM; initial $O_2$ concentration, 250 µM; and where indicated, catalase, 500 U $ml^{-1}$, or superoxide dismutase, 100 U $ml^{-1}$; irradiance, 2200 µmol photons $m^{-2}\ s^{-1}$.

The ET from PSI cofactors to $O_2$ results in $O_2^{\bullet-}$ generation supporting the stoichiometry 1 $e^-$ : 1 $O_2\downarrow$:

$$2\ e^- + 2\ O_2\downarrow \rightarrow 2\ O_2^{\bullet-} \quad (1)$$

The superoxide radical further reacts with either another $O_2^{\bullet-}$ *via* spontaneous dismutation or with Asc molecule, which present in suspension in excess as a part of the electron donor pair to $P_{700}^+$. When all $O_2^{\bullet-}$ dismutate, the stoichiometry is 2 $e^-$ : 1 $O_2\downarrow$ due to a release of 1 molecule of $O_2$ per 2 $e^-$:

$$2\ O_2^{\bullet-} + 2\ H^+ \rightarrow 1\ H_2O_2 + 1\ O_2\uparrow \quad (2)$$

When all $O_2^{\bullet-}$ react with Asc, the stoichiometry is 1 $e^-$ : 1 $O_2\downarrow$:

$$2\ O_2^{\bullet-} + 2\ AscH^- + 2\ H^+ \rightarrow 2\ H_2O_2 + 2\ Asc^{\bullet-} \quad (3)$$

Therefore, $O_2$ consumption rate is equal to the rate of ET through PSI only if the reaction (3) efficiently suppresses the reaction (2). The latter in our experiments was confirmed by addition of superoxide dismutase (SOD) enhancing the reaction (2) and fully suppressing the reaction (3). SOD led to a two-fold decrease in the rate of $O_2$ uptake, indicating that, in the absence of SOD, all $O_2^{\bullet-}$ produced indeed reacts with Asc avoiding dismutation.

The same stoichiometry is valid in the presence of MV, since a single PS I turnover also results in $O_2^{\bullet-}$ generation *via* the reaction of the radical form of MV with $O_2$.

Similarly to SOD, catalase also changes the stoichiometry to 2 $e^-$ : 1 $O_2\downarrow$ due to a release of 1 molecule of $O_2$ from $H_2O_2$ decomposition:

$$2\ H_2O_2 \rightarrow 2\ H_2O + 1\ O_2\uparrow \quad (4)$$

**Table S3.** Effect of Asc on the rate of $O_2$ uptake in a suspension of intact PSI complexes

| | Ascorbate | Rate of $O_2$ uptake, µmol $O_2$ (mg Chl h)$^{-1}$ |
|---|---|---|
| No additions | No | 0.19 |
| | 5 mM | 4.68 |
| Chemical quencher of $^1O_2$ (5 mM CAT1-H) | No | 0.48 |
| | 5 mM | 4.68 |

Mature wild type PSI complexes, 70 µg Chl ml$^{-1}$; initial $O_2$ concentration, 250 µM; irradiance, 750 µmol photons m$^{-2}$ s$^{-1}$.

In addition to $O_2^{\bullet -}$, singlet $O_2$ ($^1O_2$) can also contribute to $O_2$ uptake. Although PSI has never been confirmed as a source of $^1O_2$, the possibility of such a reaction should still be considered. This arises from the fact that, under steady-state illumination of PSI with molecular $O_2$ as the only acceptor from ET cofactors, the contribution of charge recombination might be high and this could lead to formation of $^3P_{700}$, which is quenched by oxygen molecules (triplet in the ground state), producing a highly reactive $^1O_2$. If it occurs, this reaction would contribute to the measured $O_2$ uptake since Asc acts as an efficient chemical quencher of $^1O_2$, reducing it to $O_2^{\bullet -}$and then to $H_2O_2$. To verify a possible $^1O_2$ contribution to the measured rate of $O_2$ uptake, we illuminated PSI in the absence of either Pc or Asc as electron donors, providing a higher yield of charge recombination. Asc as a chemical quencher of both $^1O_2$ and $O_2^{\bullet -}$ was replaced with a cyclic hydroxylamine 1-Hydroxy-2,2,6,6-tetramethylpiperidin-4-yl-trimethylammonium (CAT1-H), which reduces both $^1O_2$ to $O_2^{\bullet -}$ and $O_2^{\bullet -}$ to $H_2O_2$, but, in contrast to Asc, does not reduce $P_{700}^+$ (Kozuleva et al., 2015a). There was very little $O_2$ uptake in this experiment.

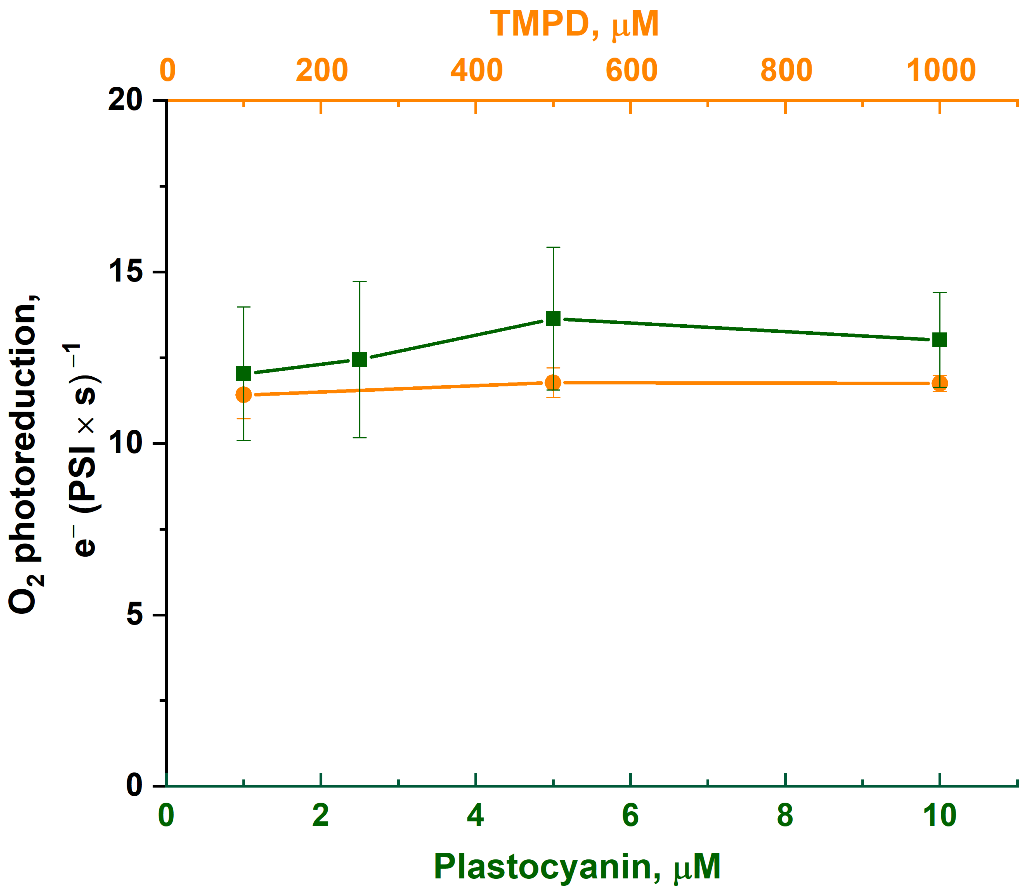


**Fig.S1.** Effect of concentration of plastocyanin (green) or TMPD (orange) as $P_{700}^{+}$ electron donor on the $O_2$ photoreduction rate. The experiments conditions were: Mature wild type PSI complexes, 15 nM; Asc, 10 mM; initial $O_2$ concentration, 250 µM; irradiance, 2200 µmol photons $m^{-2}$ $s^{-1}$.

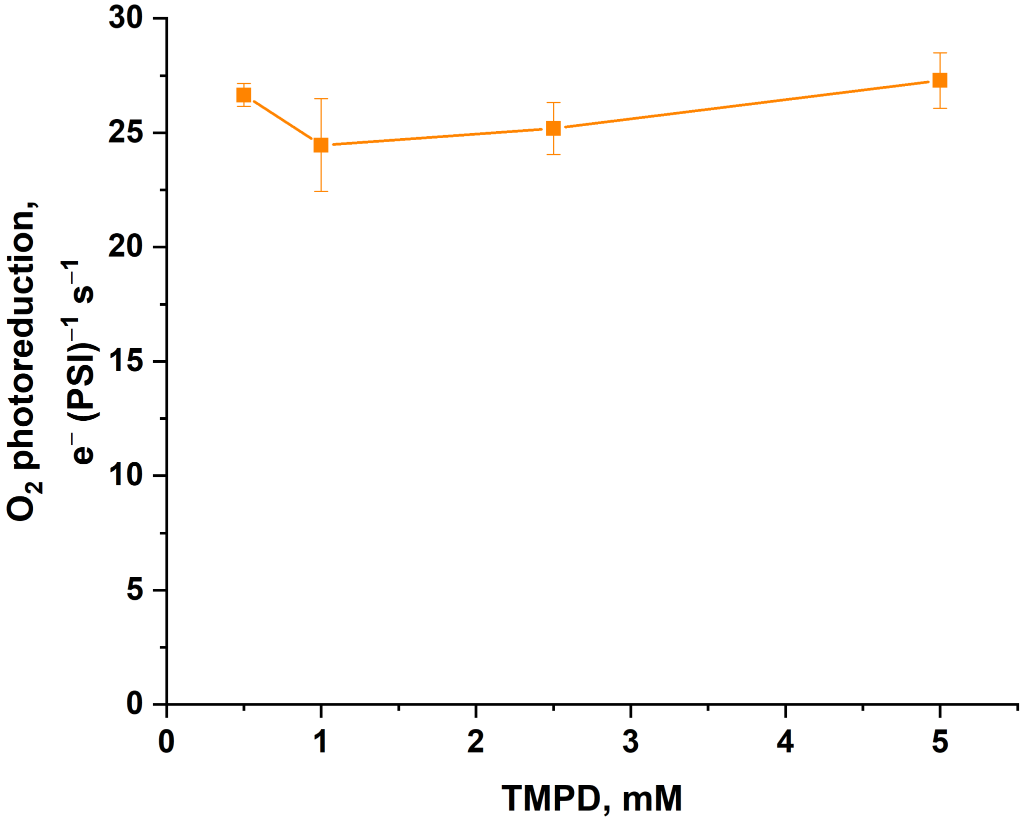


**Fig.S2.** Effect of TMPD concentration on the $O_2$ photoreduction rate in the $A_1$-core complexes. The experiments conditions were: $A_1$-core complexes from wild type, 10 nM; Asc, 10 mM; irradiance, 2200 µmol photons $m^{-2}$ $s^{-1}$; initial $O_2$ concentration, 250 µM.

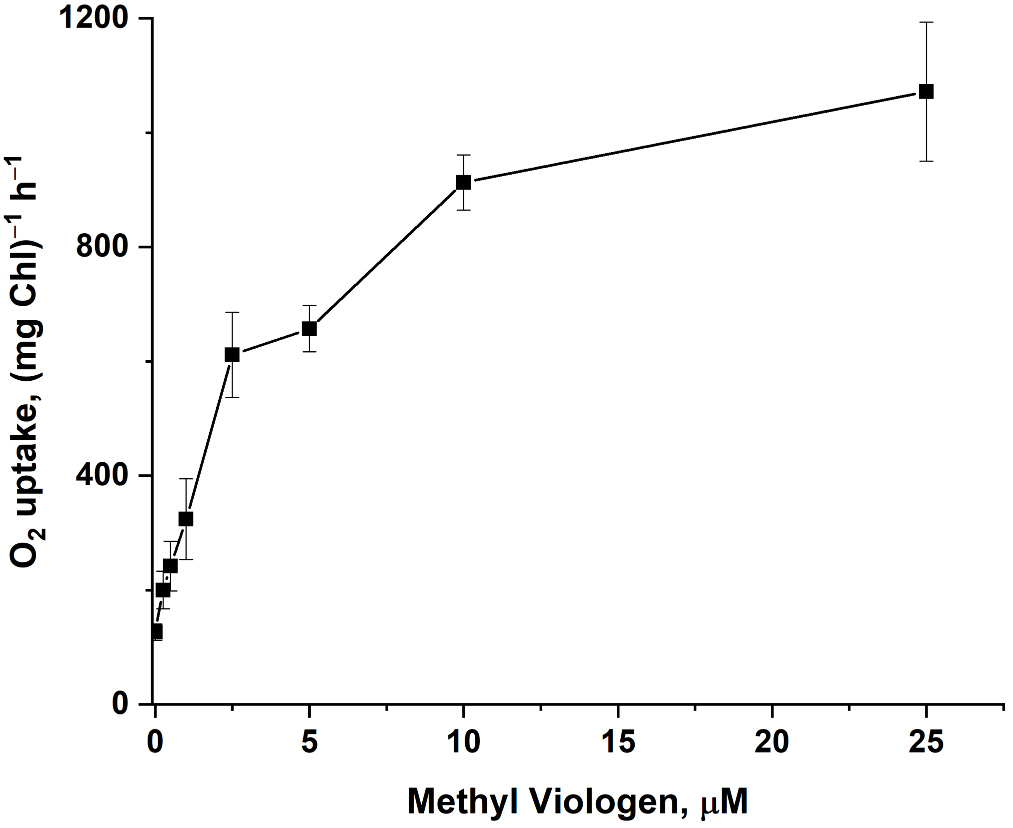


**Fig. S3.** Rate of light-induced $O_2$ uptake in a PSI suspension as a function of MV concentration. The experiments conditions were: Mature wild type PSI complexes, 10 nM; Pc, 5 μM; Asc, 10 mM; initial $O_2$ concentration, 276 μM; irradiance, 900 μmol photons $m^{-2}$ $s^{-1}$.

### Material and Methods:

#### Determination of $K_m$(MV)

The apparent $K_m$(MV) was calculated from the analysis of dependence of the $O_2$ uptake rate on MV concentration, using software package Origin 2017. To calculate the apparent $K_m$(MV), the rate of direct $O_2$ photoreduction by PSI in the absence of MV was subtracted from the rates observed in the presence of MV (Petrova et al., 2018).

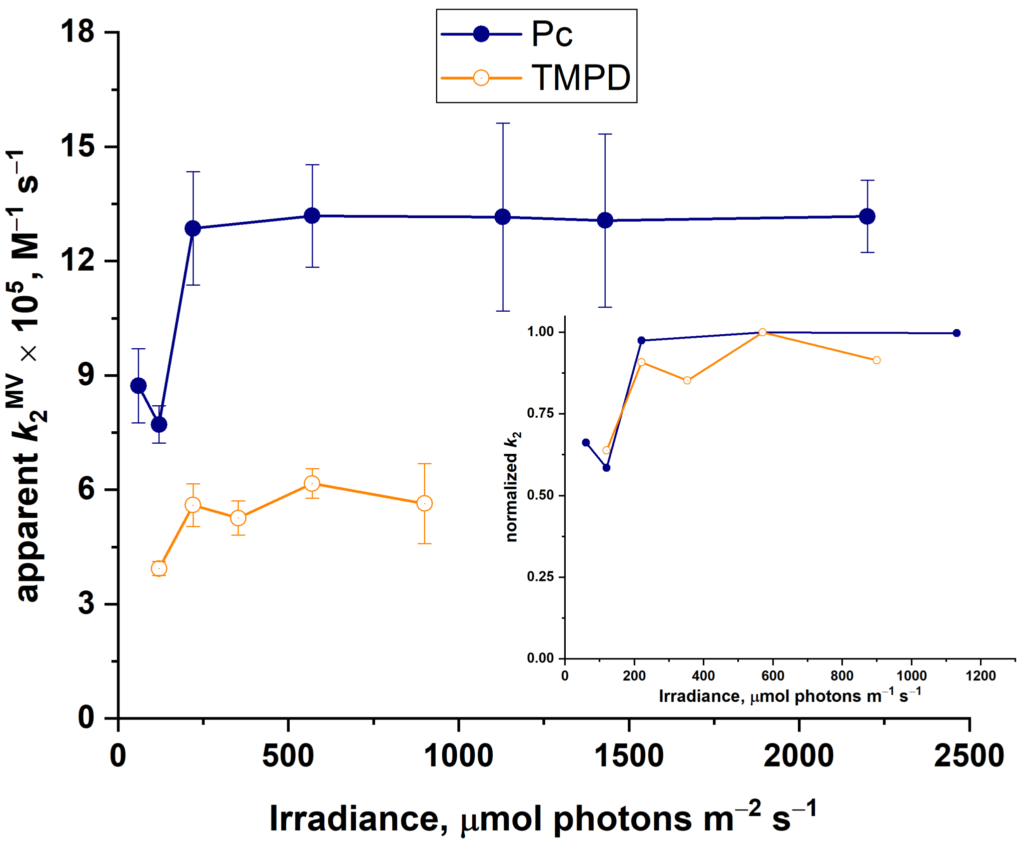


**Fig. S4.** Effect of irradiance on the apparent $k_2^{MV}$ in the presence of Pc (5 μM; dark blue lines) or TMPD (1 mM; orange line) as electron donor to $P_{700}^+$. Insert: normalized to the maximal $k_2^{MV}$ values for (B). The experiments conditions were: Mature wild type PSI complexes, 10 nM; MV, 50 μM; Asc, 10 mM; catalase, 500 U $ml^{-1}$; initial $O_2$ concentration, 10-30 μM.

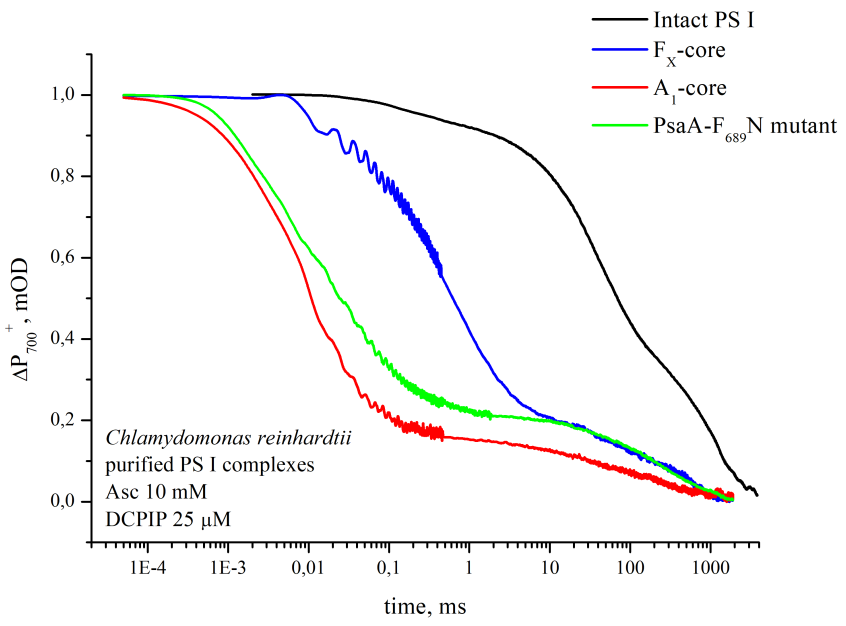


**Fig. S5.** Charge recombination in the intact PSI, $F_X$-core, and $A_1$-core complexes from WT and intact mature PSI from the PsaA-$F_{689}$N mutant.

### Material and Methods:

**Measurement of Charge recombination kinetics**. Flash-induced $P_{700}^+$ reduction kinetics were monitored at 820 nm using a laboratory-built spectrophotometer where the source of the measuring light was a laser diode, while saturating light flashes were produced by a frequency-doubled Quantel Nd:YAG laser (wavelength, 532 nm; pulse half-width, 12 ns; flash intensity, 20 mJ). Experiments were performed in a standard 1-cm optical path glass cuvette containing 50 mM HEPES-NaOH buffer (pH 7.5), 0.03% w/v β-DDM, and purified PSI complexes at a Chl concentration 100 μg $ml^{-1}$. Electron donors added were 15 μM 2,3-dichlorophenolindophenol and 10 mM ascorbate. Generally, 16 signals with 10 s intervals were averaged.

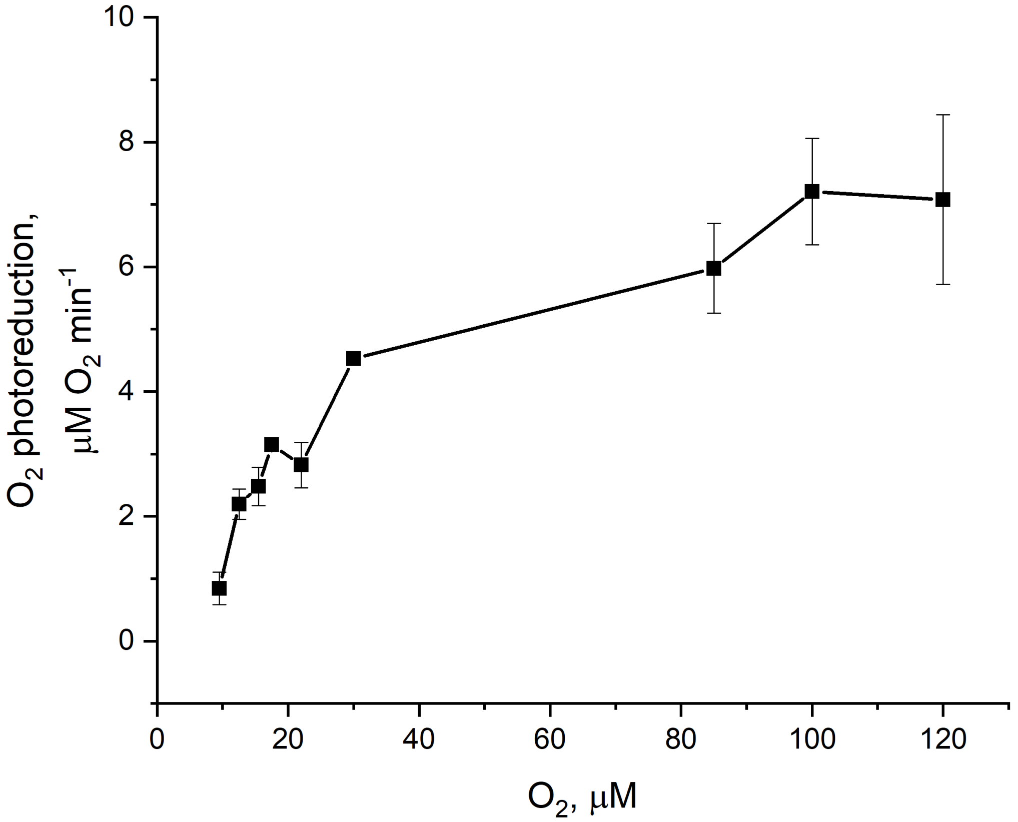


**Fig. S6.** Rate of light-induced $O_2$ uptake in a PSI suspension as a function of $O_2$ concentration**.** The experiments conditions were: Mature wild type PSI complexes, 15 nM; Pc, 5 µM; Asc, 10 mM; irradiance, 2200 µmol photons $m^{-2}$ $s^{-1}$.

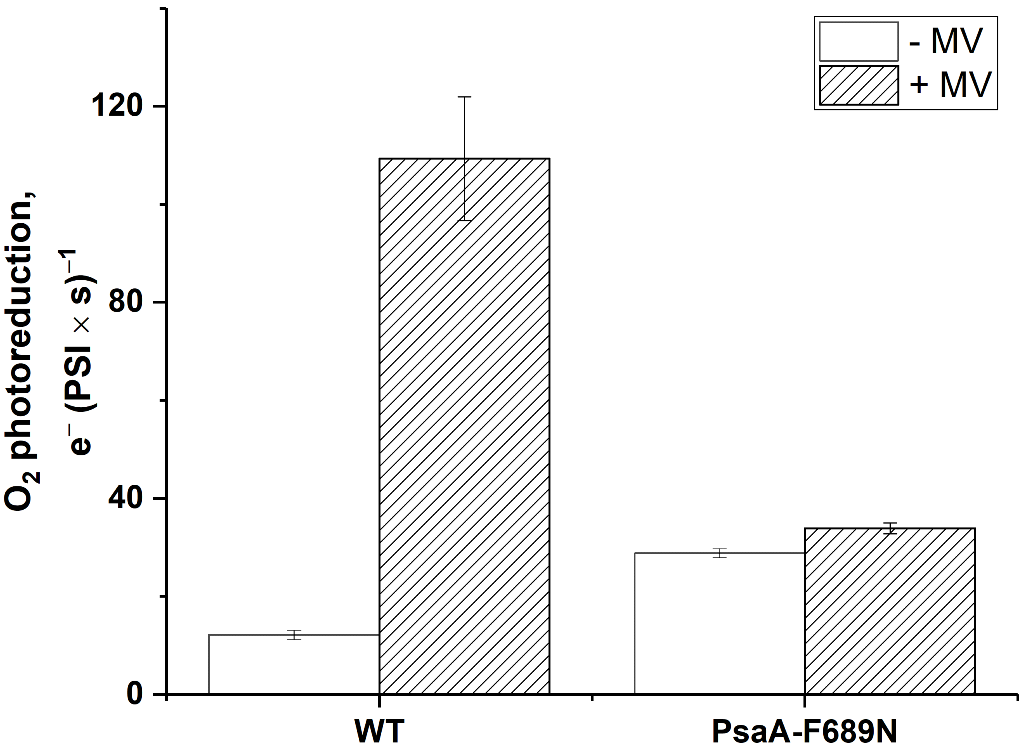


**Fig. S7.** Effect of MV addition on the rate of $O_2$ photoreduction in PSI complexes from WT and PsaA-$F_{689}$N mutant. The experiments conditions were: Mature PSI complexes from wild type or PsaA-$F_{689}$N, 10 nM; Asc, 10 mM; Pc, 5 µM; catalase, 500 U $ml^{-1}$; initial $O_2$ concentration, 250 µM; irradiance, 2200 µmol photons $m^{-2}$ $s^{-1}$.

Appendix

**Calculation of the apparent rate constant of $O_2$ photoreduction by PSI**

The reduction of $O_2$ by PSI ($V_{O2}$) is a second order reaction:

$$V_{O2} = k_2 \times [PSI] \times [O_2] \quad (1)$$

Since the concentration of PSI, which was equal to concentration of $P_{700}^{+}$, did not change, $k_2 \times$ [PSI] was expressed as a pseudo-first order rate constant ($k_1'$):

$$V_{O2} = k_1' \times [O_2] \quad (2)$$

The $k_1'$ was calculated as:

$$k_1' = \frac{1}{t} \times \ln \frac{[O2\,]^0}{[O2]^t} \quad (3)$$

where $t$ represents the duration of illumination (s), $[O_2]^0$ and $[O_2]^t$ represent the concentrations of oxygen at the onset and at t timepoints during the illumination, respectively. It is necessary to correct $[O_2]^t$ to account for non-specific $O_2$ concentration changes occurring in the dark and in the light. It is also necessary to correct for the addition of catalase, which decomposes $H_2O_2$ to release one $O_2$ per two $H_2O_2$ molecules produced *via* $O_2$ photoreduction by PSI (Table S2). Therefore the $[O_2]^t$ values in eq. 3 were based on calculating light-induced $V_{O2}$:

$$[O_2]^t = [O_2]^0 - t \times 2 \times V_{O2} \quad (4)$$
